# New kinship and *F*_ST_ estimates reveal higher levels of differentiation in the global human population

**DOI:** 10.1101/653279

**Authors:** Alejandro Ochoa, John D. Storey

## Abstract

Kinship coefficients and *F*_ST_, which measure genetic relatedness and the overall population structure, respectively, have important biomedical applications. However, existing estimators are only accurate under restrictive conditions that most natural population structures do not satisfy. We recently derived new kinship and *F*_ST_ estimators for arbitrary population structures [1, 2]. Our estimates on human datasets reveal a complex population structure driven by founder effects due to dispersal from Africa and admixture. Notably, our new approach estimates larger *F*_ST_ values of 26% for native worldwide human populations and 23% for admixed Hispanic individuals, whereas the existing approach estimates 9.8% and 2.6%, respectively. While previous work correctly measured *F*_ST_ between subpopulation pairs, our generalized *F*_ST_ measures genetic distances among all individuals and their most recent common ancestor (MRCA) population, revealing that genetic differentiation is greater than previously appreciated. This analysis demonstrates that estimating kinship and *F*_ST_ under more realistic assumptions is important for modern population genetic analysis.

---

Kinship coefficients and *F*_ST_ are defined as probabilities of identity-by-descent [3–5]. Kinship matrices are crucial for accurate inference under population structure in many important biomedical applications, including genome-wide association studies [6–13] and heritability estimation [14, 15]. However, the most commonly-used standard kinship estimator [9, 10, 13–19] is accurate only in the absence of population structure [2, 20]. Likewise, current *F*_ST_ estimators assume that individuals are partitioned into statistically-independent subpopulations [4, 5, 21–23], which does not hold for human and other complex population structures. The human genetic population structure is remarkably complex, shaped by geography and population bottlenecks in migrations out of Africa [24–34] and admixture events [35–39]. We use human data to illustrate the improvements provided by our new approach.

## Models and methods

Our new kinship and *F*_ST_ estimators were derived assuming arbitrary population structures, and they yield nearly unbiased estimates [2]. Suppose there are *n* individuals genotyped at *m* biallelic autosomal loci, such as SNPs. Our kinship estimator 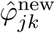 is given by

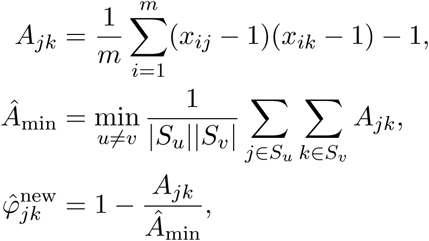

where the genotypes *x*_*ij*_ ∈ {0, 1, 2} count the number of reference alleles at locus *i* for individual *j*. For simplicity, here *Â*_min_ uses a partition of individuals into subpopulations *S*_*u*_ for *u* ∈ {1, …, *K*} used solely to estimate the minimum kinship, which is set to zero (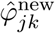 has individual-level resolution; the general framework does not need subpopulations [2]). Under our model E[*A*_*jk*_] = (*φ*_*jk*_ −1)*v* contains the desired kinship coefficient *φ*_*jk*_ and a nuisance parameter *v* shared by all individuals. Assuming zero kinship across the two least related individuals, E[*Â*_min_] ≈ min_*j,k*_ E[*A*_*jk*_] = −*v* yields the nuisance parameter, enabling consistent kinship estimates: 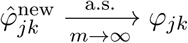.

We compare to the widely-used standard kinship estimator

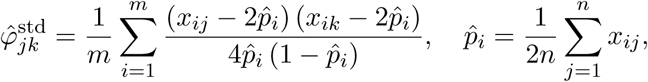

which has a complex bias non-linear in *φ*_*jk*_ in structured populations [2, 20]. The limit of 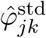 as the number of loci *m* → ∞ is well-approximated by

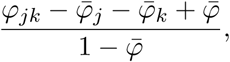

where 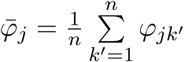 is the mean kinship of individual *j* with all others and 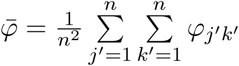 is the overall mean kinship in the data [2]. This estimator is widely-used in approaches for structured populations, including genetic association studies and heritability estimation [9, 10, 13–19].

The original *F*_ST_ measures inbreeding in a subpopulation relative to an ancestral population [4], excluding local inbreeding if present [5]. Existing approaches estimate the mean *F*_ST_ between two or more independent subpopulations relative to their MRCA population [21, 23, 40], but have a downward bias otherwise [2]. In our new approach, inbreeding coefficients are estimated from kinship (measured from the MRCA population) using 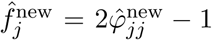 and the generalized *F*_ST_ is estimated using 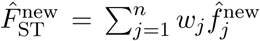 (valid for locally-outbred individuals [1]), where *w*_*j*_ are weights to account for geographically-imbalanced sample sizes. Our generalized *F*_ST_ does not require subpopulations, is the first to be applicable to arbitrary population structures [1], and our estimator is accurate in this setting [2]. We compare to the existing *F*_ST_ estimators Weir-Cockerham [21], HudsonK (for two subpopulations [23] generalized in [2]), and BayeScan [40]. These estimators assume independent subpopulations and homogeneous inbreeding within subpopulations, which causes downward biases in more complex population structures and implicitly admit negative kinship coefficients [2]. The classical *F*_ST_ interpretation—the proportion of variance explained by differences between subpopulation pairs—is not appropriate when subpopulations are not independent, and is not clearly defined in the absence of obvious subpopulations (such as for admixed individuals). Instead, our generalized *F*_ST_ measures the genetic drift of individuals from the MRCA population, which ensures valid underlying kinship coefficients [1].

## Results

We first analyze the Human Origins datasets of native populations [41–43], which consists of **2922 individuals** from **243 sub-subpopulations** grouped into **11 subpopulations** (Supplementary Information). Sub-subpopulation abbreviations are defined in [41–43], while the subpopulation labels are defined in Fig. S1 (Supplementary Information). If these subpopulations were independent and internally unstructured, as assumed by existing *F*_ST_ estimators, the kinship matrix would have zero values between subpopulations and equal kinship within subpopulations (Fig. 1A). Instead, our approach reveals substantial kinship between subpopulations and heterogeneity within subpopulations (Fig. 1C).

**Figure 1:**
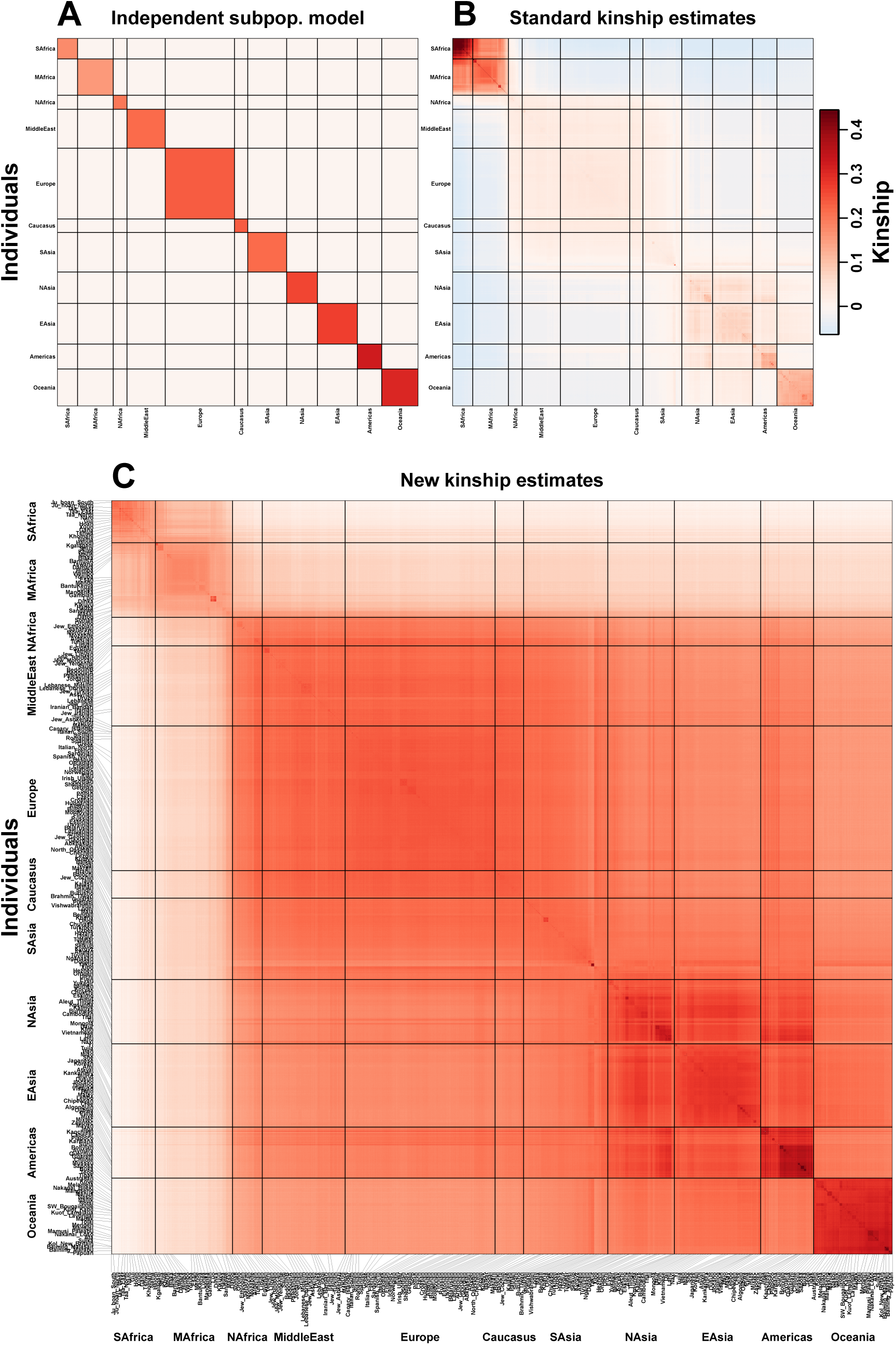
Population-wide kinship estimates in Human Origins. As a visual aid, individuals are arranged into a hierarchy with subpopulations (rough continental clusters, i.e., `SAfrica`) and sub-subpopulations (locations potentially separated by ethnicity or religion, i.e., `Lebanese_Christian`). However, we estimate individual-level kinship without using this hierarchy. Color corresponds to kinship (*φ*_*jk*_) for every pair of individuals *j* (x-axis) and *k* (y-axis) and inbreeding coef. (*f*_*j*_) along the diagonal. **A.** Kinship matrix assumed by the independent sub-populations model prevalent in *F*_ST_ estimation: fixed *φ*_*jk*_ within subpopulations, *φ*_*jk*_ = 0 between subpopulations. **B.** Biased standard kinship estimates. The overall downward bias causes many negative estimates (blue) and strong distortions across the matrix (incorrectly assigns highest kinship within `SAfrica`-`MAfrica`), as predicted by our theory. Also note comparable kinship estimates between each of `MAfrica`, `Europe`, and `EAsia`, which contradicts the African origins model. **C.** New kinship estimates. Our new estimates reveal substantial kinship between subpopulations and heterogeneity within subpopulations. For an improved dynamic range, all displayed *f*_*j*_, *φ*_*jk*_ values in panels B and C were capped at the 99 percentile of the estimated *f*_*j*_ values of panel C (full *f*_*j*_ distribution in Fig. 3). Additionally, panel B was capped below to the 1 percentile of its distribution.

The kinship matrix of Fig. 1C can be interpreted under the African origins serial founder model, as follows. Recall that a population size reduction (bottleneck) increases kinship and *F*_ST_ relative to the ancestral population [3–5]. The first population split occured roughly between individuals from Sub-Saharan African (KhoeSan-speaking hunter-gatherers (`SAfrica`) and Bantu-speakers and other agro-pastoralists (`MAfrica`)) and individuals outside of Sub-Saharan Africa. This split resulted in bottlenecks that increased kinship in each side relative to the ancestral value (which equals the kinship between the two subpopulations). The next split was roughly between West Eurasians and the rest, again increasing kinship within each side. Among West Eurasians, kinship is higher within `Europe`, reflecting another bottleneck. `Americas` (Native Americans) and `Oceania` have the highest kinship values within, reflecting further bottlenecks in their trek out of Africa. Note that the European admixture in `Americas` (calculated in Supplementary Information) is evident in individuals with lower kinship relative to other `Americas` individuals and greater kinship with `Europe` (Fig. 1C). Overall, our observations are coherent with previous work [27, 30, 31, 35], but our approach is the first to use a nearly unbiased estimator of kinship coefficients under assumptions aligned with the data. Our approach accurately estimates kinship at individual-level resolution and successfully uncovers a complex population structure where individuals may be related to each other in arbitrary ways.

The MRCA population of living humans is estimated to have existed in Africa 100-200K years ago [26, 27, 34], which first split into the ancestral KhoeSan population (who speak so-called “click” languages of the Kx’a, Tuu, and Khoe families, grouped into `SAfrica`) and the rest [26, 27, 31, 32, 34]. This MRCA population excludes ancestry from the Neanderthal and Denisovan introgressions [36, 37], but their limited contribution makes it a reasonable approximation. In our estimates, the minimum per-sub-subpopulations mean kinship is between `Ju_Hoan_North` (`SAfrica`) and `Kol_New_Britain` (`Oceania`). Moreover, the 2114 pairs with the smallest kinship values all consisted of pairs where one sub-subpopulation was from `SAfrica` (most commonly `Ju_Hoan_North` and `Ju_Hoan_South`) and the other was from outside of `SAfrica` and `MAfrica`. Therefore, we infer the first population split to have been between the ancestral KhoeSan population (`SAfrica`) and the rest, agreeing with previous work using independent mtDNA [26], Y chromosome [34], and microsatellite data [27, 32] (not used by our approach), as well as SNPs [31]. High kinship between `SAfrica` and `MAfrica` (Fig. 1C) suggests recent admixture [32] or an isolation-by-distance structure [44].

The diagonal of the kinship matrix of Fig. 1C contains inbreeding coefficients *f*_*j*_, which are individual-specific *F*_ST_ values (for locally-outbred individuals, which most humans are). Every individual is differentiated (first percentile *f*_*j*_ = 0.149, where the zero value would correspond to the *f*_*j*_ of the child of the two most unrelated individuals in the data), and differentiation increases with distance from southern Africa (shown geographically in Fig. 2 and using distributions in Fig. 3), as expected under the African origins model and agreeing with previous work [26, 27, 29–34]. Remarkably, our estimated *F*_ST_ of 0.260 is substantially larger than estimates around 0.098 from existing approaches (Fig. 3) and previous measurements based on *F*_ST_ [30, 45] or related variance component models [31, 46, 47] — except for some AMOVA *ϕ*_ST_ estimates [48] (pairwise *F*_ST_ estimates [23, 49–52] are not generally comparable to our estimate). Existing approaches underestimate *F*_ST_ because they assume zero kinship between subpopulations, clearly incorrect as seen in Fig. 1C, whereas our new approach models arbitrary kinship between individuals and leverages kinship to estimate *F*_ST_.

**Figure 2:**
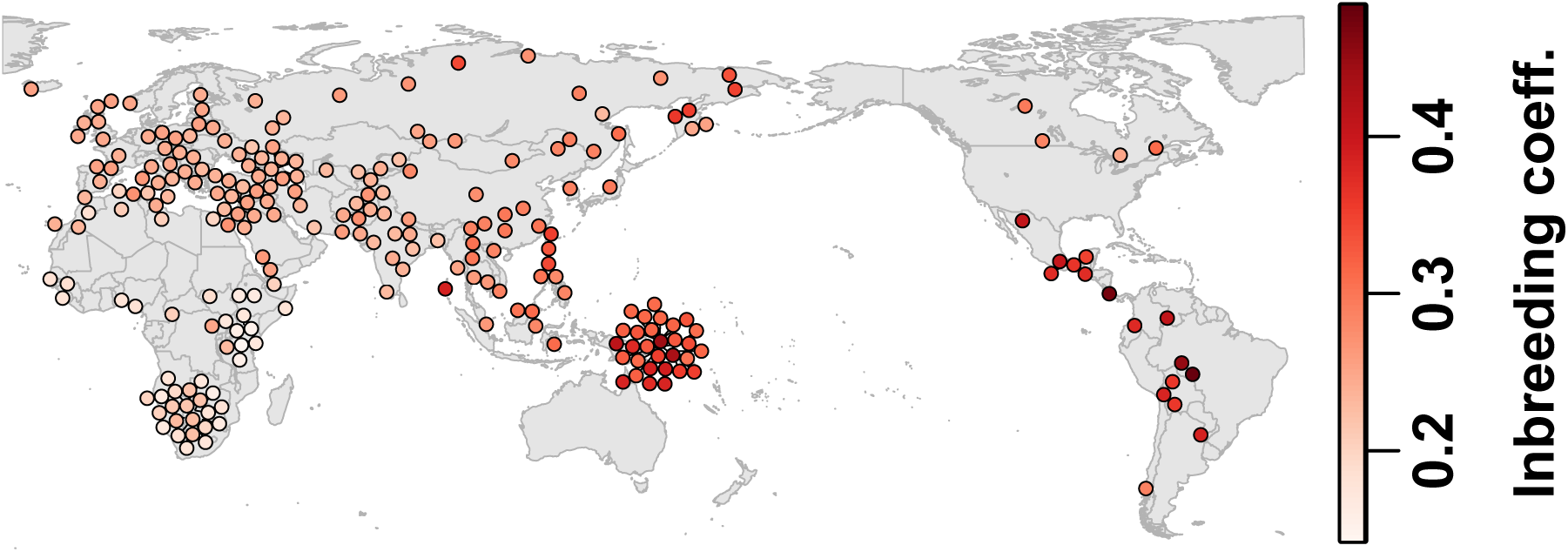
Geographical distribution of population-level inbreeding. Colors in circles denote the mean individual inbreeding *f*_*j*_ within each Human Origins sub-subpopulation. These mean *f*_*j*_ values increase smoothly with distance from southern Africa, as expected under the African origins serial founder model.

**Figure 3:**
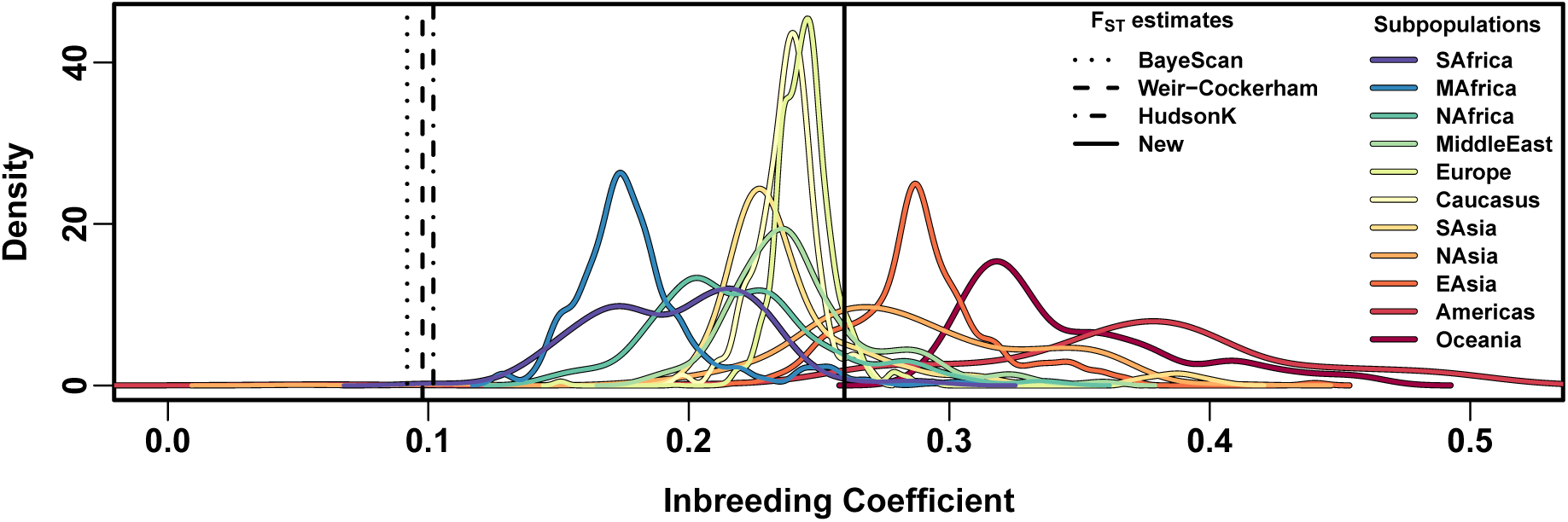
Inbreeding and *F*_ST_ estimates in Human Origins. Our new approach yields individual *f*_*j*_ inbreeding values and a generalized *F*_ST_ estimate, which is their weighted mean. Per-subpopulation *f*_*j*_ distributions show increasing values with distance from Africa (where densities are estimated with equal sub-subpopulation weights). Weir-Cockerham, HudsonK, and BayeScan assumed the *K* = 11 subpopulations are independent (Fig. 1A), which causes downward bias.

The popular standard kinship estimator [9, 10, 13–19] has a nonlinear bias in structured populations [2, 20]. Standard kinship matrix estimates have abundant negative values and strong distortions (Fig. 1B *versus* our estimates in Fig. 1C, direct comparison in Fig. S2A). These estimates disagree with the African origins model, assigning greater kinship within `SAfrica`-`MAfrica` than to any other subpopulation, and comparable kinship between `Europe`, `EAsia` and `MAfrica` (incorrect since `MAfrica` split first [23]). The biases in the standard kinship and existing *F*_ST_ estimators are both fundamentally due to assuming that most kinship values are zero, and explicitly or implicitly admit negative kinship values [2, 22]. Our new kinship estimator is developed for arbitrary population structures and yields more interpretable estimates in human data.

Hispanic Latin Americans have a complex population structure, being recently admixed from European (`EUR`), Native American (`AMR`), and Sub-Saharan African (`AFR`) populations [53–57]. Here we show that Hispanics in the 1000 Genomes Project (TGP) do not have discrete subpopulations, so the classical *F*_ST_ definition does not apply. Using our approach, we estimate the kinship matrix of the 347 TGP Hispanic individuals (`PUR`: Puerto Rican; `CLM`: Colombian; `PEL`: Peruvian; `MXL`: Mexican-American; standard kinship in Fig. 4A, our new approach in Fig. 4B). We also estimate individual-specific admixture proportions of `EUR`, `AMR`, and `AFR` ancestry (Fig. 4C), detailed in Supplementary Information. We confirm previous observations that relatedness in Hispanics varies along a continuum driven by admixture [53–57]. In particular, since differentiation increases from `AFR` to `EUR` to `AMR` (Fig. 3), the greatest kinship is between individuals with higher `AMR` ancestry, and the lowest kinship is between individuals with higher `AFR` ancestry (Fig. 4B and C). Standard kinship estimates are also biased and distorted in Hispanics (Fig. 4A and Fig. S2B) and lack the interpretability of our estimates. Lastly, our approach estimates *F*_ST_ to be 0.233, which is comparable to *f*_*j*_ estimates for `MAfrica` and `Europe` in Human Origins (Fig. 3). In contrast, Weir-Cockerham estimates an unrealistically small *F*_ST_ of 0.0260, which is downwardly biased because it requires subpopulation labels (we used sampling locations: `PUR`, `CLM`, `PEL`, `MXL`; see the thin colored bar outside each kinship matrix in Fig. 4A and B), which erases the considerable substructure within subpopulations and models the large kinship values between individuals of different subpopulations as zero. We emphasize that existing *F*_ST_ approaches were designed for and require non-overlapping, independently evolving subpopulations, so they do not apply to individuals with variable admixture proportions such as Hispanics (as shown here) and African Americans [32, 58]. Our results are not a critique of those important advances, but a demonstration that modern data requires new estimators. Our approach overcomes these challenges by estimating kinship without assuming subpopulations (Supplementary Information) and estimating *F*_ST_ from these individual-specific parameters.

**Figure 4:**
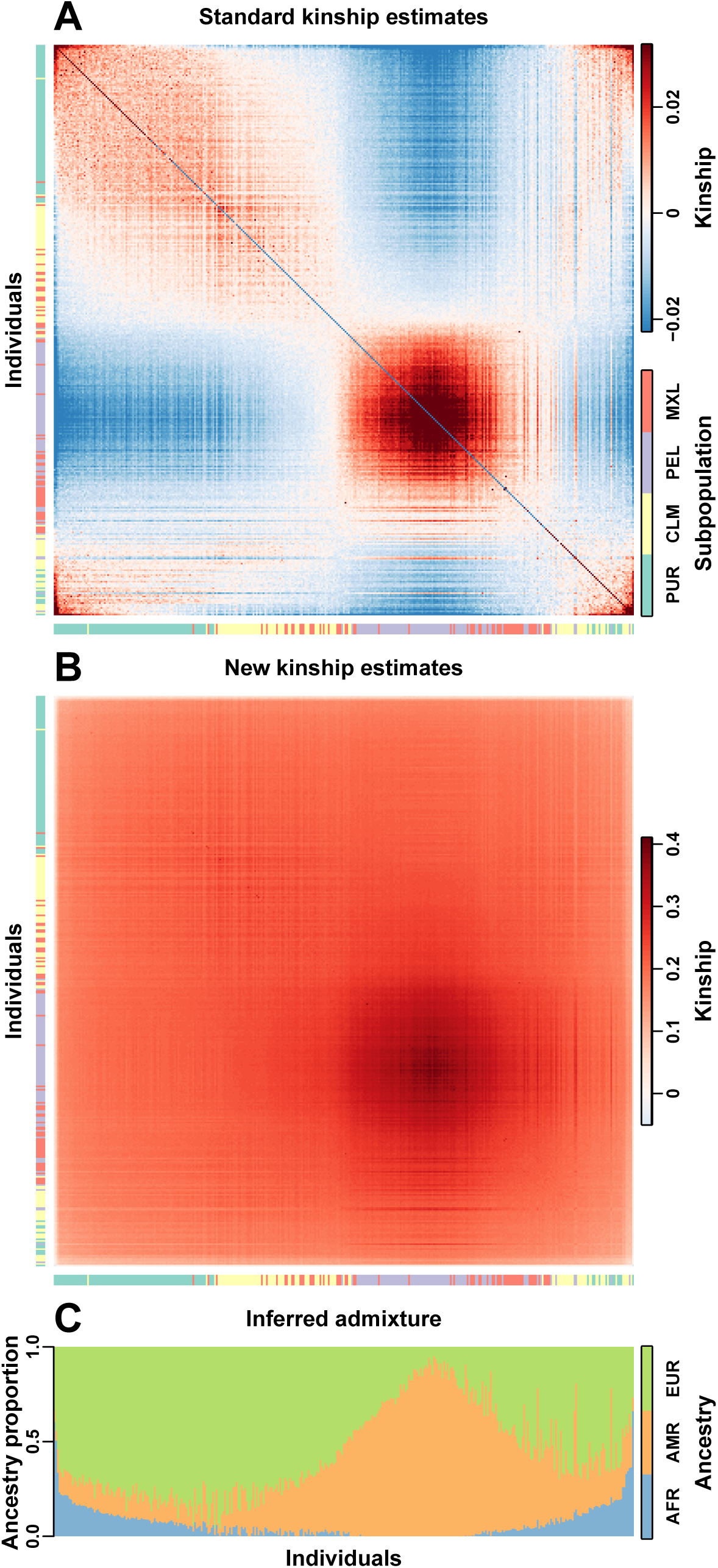
Kinship of Hispanics in 1000 Genomes. The colors in the kinship heatmaps correspond to *φ*_*jk*_ kinship values for every pair of individuals *j* (x-axis) and *k* (y-axis), and *f*_*j*_ along the diagonal. The color bars outside kinship matrices mark the subpopulation (sampling location) of every individual, which is poorly correlated with kinship (panel B) or admixture proportions (panel C). **A**: Biased standard kinship estimates. Note the overall downward bias causes negative estimates (blue), discontinuities between *f*_*j*_ (diagonal) and *φ*_*jk*_ (off-diagonal), and distortions (e.g., relatively higher kinship between individuals with higher `AFR` ancestry). Displayed *φ*_*jk*_, *f*_*j*_ values are capped to the 1 and 99 percentiles of their distribution. **B**: New kinship estimates reveal a smooth population structure without discrete sub-populations, and much greater kinship values (note the different color scales for panels A and B). Individuals were ordered using seriation, which places low *φ*_*jk*_ away from the diagonal [59, 60]. **C**: Admixture proportions of every individual for Sub-Saharan African (`AFR`), European (`EUR`), and Native American (`AMR`) ancestry (calculated in Supplementary Information).

## Conclusion

Our analysis of the Human Origins and 1000 Genomes Project datasets reveals a complex population structure with predominantly non-zero kinship values that vary along a continuum. Our new approach does not artificially partition individuals into subpopulations, which enables us to capture population structure with individual-level resolution and for the first time yields accurate population-level kinship and *F*_ST_ estimates for world-wide human SNP data. New approaches that make minimal assumptions about relatedness and structure are necessary for many biomedical applications—including genetic association studies in multiethnic panels and admixed individuals—and our new framework provides the foundation that enables this goal.

## Data and Software

This approach is implemented in the R package `popkin` available online (https://cran.r-project.org/package=popkin and https://github.com/StoreyLab/popkin). Public data and code reproducing these analyses are available at https://github.com/StoreyLab/human-differentiation-manuscript.

## Acknowledgments

This research was supported in part by NIH grant R01 HG006448.

## S1 Human Origins

### S1.1 Data Processing

The Human Origins data described in the main text is merged from several sources [41–43] and processed as follows. Both Original and Pacific in Table S1 refer to the full datasets (public and non-public portions) described in [41–43] after removing non-autosomal loci and excluding ancient individuals. The Final dataset described in the main text is the union of individuals in Original and Pacific and the intersection of loci, after additional filters described below. This dataset was processed using the plink2 software [61].

**Table S1:**
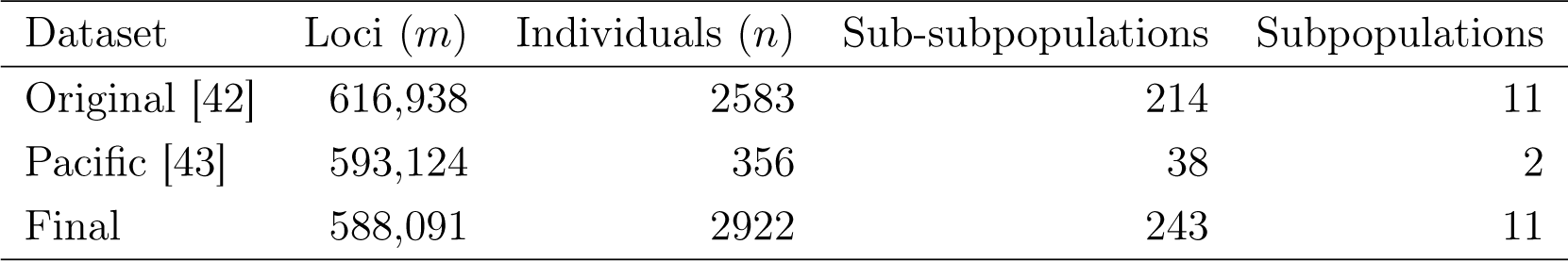
Overview of Human Origins datasets and filters.

| Dataset | Loci ($m$) | Individuals ($n$) | Sub-subpopulations | Subpopulations |
| --- | --- | --- | --- | --- |
| Original [42] | 616,938 | 2583 | 214 | 11 |
| Pacific [43] | 593,124 | 356 | 38 | 2 |
| Final | 588,091 | 2922 | 243 | 11 |

We obtained the full (public and non-public) Human Origins data presented in [41–43] by contacting the authors and agreeing to their usage restrictions. The public subset of these data is available at https://reich.hms.harvard.edu/datasets. The Original dataset described in [41, 42] was obtained as a single, merged dataset. The Pacific dataset described in [43] was obtained as a separate dataset. Geographical coordinates for these individuals were obtained from supplementary data [41–43].

Both Original and Pacific were genotyped on the same microarray platform, but a small subset of loci present in Original were removed from Pacific due to more stringent quality checks (David Reich, personal communication). For that reason, we merged Original and Pacific by considering the *union* of individuals but the *intersection* of loci (Table S1).

These datasets include labels that group individuals, called here *sub-subpopulations*. We removed individuals from the singleton sub-subpopulations `“Ignore_Adygei (relative_of_HGDP01382)`”, `Saami_WGA, Wayuu, Ticuna`, and `Chane`, as well as `AA` (African Americans) since they were the only non-native sub-subpopulation. Then we removed non-polymorphic loci.

We then edited a few sub-subpopulation labels, as follows. We merged all individuals from the four subgroups `GujaratiA-D` into `Gujarati`. Lastly, the label `Southwest_Bougainville` was shortened to `SW_Bougainville` in figures. The resulting 243 native sub-subpopulations were manually grouped for visual aid into 11 subpopulations by taking into account geography, population history, and our kinship estimates (Fig. S1).

**Figure S1:**
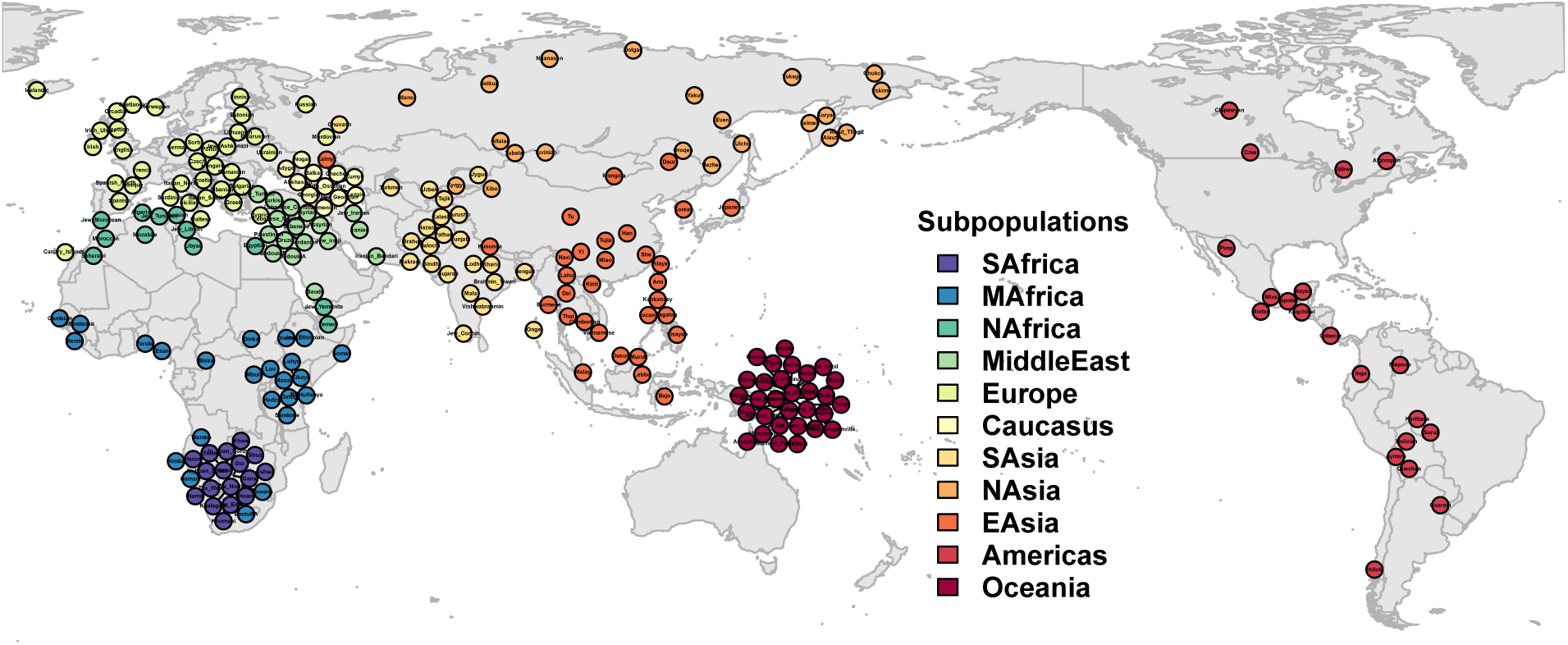
Grouping of Human Origins sub-subpopulations into *K* = 11 subpopulations. Each circle represents a sub-subpopulation, which was placed near its sampling location but nudged if necessary so circles do not overlap. The color of the circles corresponds to the subpopulation cluster we assigned. This partition into subpopulation is based on geography, history, language families, and our kinship estimates.

### S1.2 Weights for individuals

The Final dataset has a wide distribution of sub-subpopulation sample sizes, with a median sub-subpopulation size of 10 individuals, a minimum of 2 (`Canary_Islander`), and a maximum of 70 (`Yoruba`).

To calculate our generalized *F*_ST_ estimate, weights were constructed so that every subpopulation is weighed equally, and every sub-subpopulation weighs equally within each subpopulation, as follows. For every individual *j*, let *n*_*j*_ be the number of individuals in the sub-subpopulation of *j*, and *m*_*j*_ be the number of sub-subpopulations in the subpopulation of *j*. The weights used are then 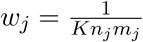, where *K* is the number of subpopulations.

### S1.3 Comparison of new and existing kinship estimates

A direct comparison of each our new kinship and inbreeding estimates on the real data to those from the standard kinship estimator are presented in Fig. S2. We found previously that bias in the standard kinship estimator varies for every pair of individuals depending on their mean kinship to everybody else in the dataset [2]. This effect is evident in our comparisons, resulting in complex patterns for the standard kinship estimates that are not linear and are not functions of the true kinship values (estimated without bias by our new estimator).

Biases in standard inbreeding coefficients (estimated as 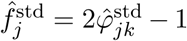; second row of Fig. S2) are considerably more extreme compared to the biases of the kinship values between different indivdiuals (first row of Fig. S2). In particular, standard inbreeding estimates often exceed 1, and in the case of Hispanics from 1000 Genomes they are negatively correlated with their true values.

### S1.4 Admixture analysis

We performed an admixture analysis to complement out analysis of kinship and *F*_ST_. We used the Admixture software [62] to infer *K* = 7 ancestry clusters from the Final dataset (see Table S1). The admixture proportions can be visualized as stacked barplots on Fig. S3, where individuals are ordered just as in the kinship matrix of the main text. These seven infered ancestry clusters correspond approximately with the 11 subpopulations we constructed based on geography and other criteria (Section S1.1) as follows (Fig. S3):

- Cluster 1 corresponds to `SAfrica.`
- Cluster 2 corresponds to `MAfrica.`
- Cluster 3 corresponds to `Europe, NAfrica, MiddleEast and Caucasus.`
- Cluster 4 corresponds to `SAsia.`
- Cluster 5 corresponds to `EAsia` and `NAsia.`
- Cluster 6 corresponds to `Americas.`
- Cluster 7 corresponds to `Oceania.`

We typically see that each ancestry cluster is concentrated in a certain geographical region, and this ancestry is also present to a lesser extent in neighboring regions and diminishes with geographical distance from its point of greatest concentration. This again argues for a complex population structure where relatedness at the population level falls on a continuum rather than taking on discrete values.

The most notable geographic discontinuities in ancestry were observed for cluster 3, which is roughly West Eurasian ancestry. Unusually high proportions of this ancestry are observed in most individuals of two sub-subpopulations of the Kamchatka Peninsula in Russia (`Aleut` and `Aleut_Tlingit` in subpopulation `NAsia`), as well as the four `Americas` sub-subpopulations from Canada (`Chipewyan, Cree, Algonquin,` and `Ojibwa`) and `Chilote` from Chile (Fig. S3). We also observed an unusual reduction of cluster 5 ancestry in `Aleut` and `Aleut_Tlingit` relative to its closest `NAsia` sub-subpopulations, and unusually low levels of cluster 6 ancestry in `Chipewyan, Cree, Algonquin, Ojibwa`, and `Chilote` relative to the rest of `Americas` sub-subpopulations. These data point to likely recent European admixture in those individuals.

**Figure S2:**
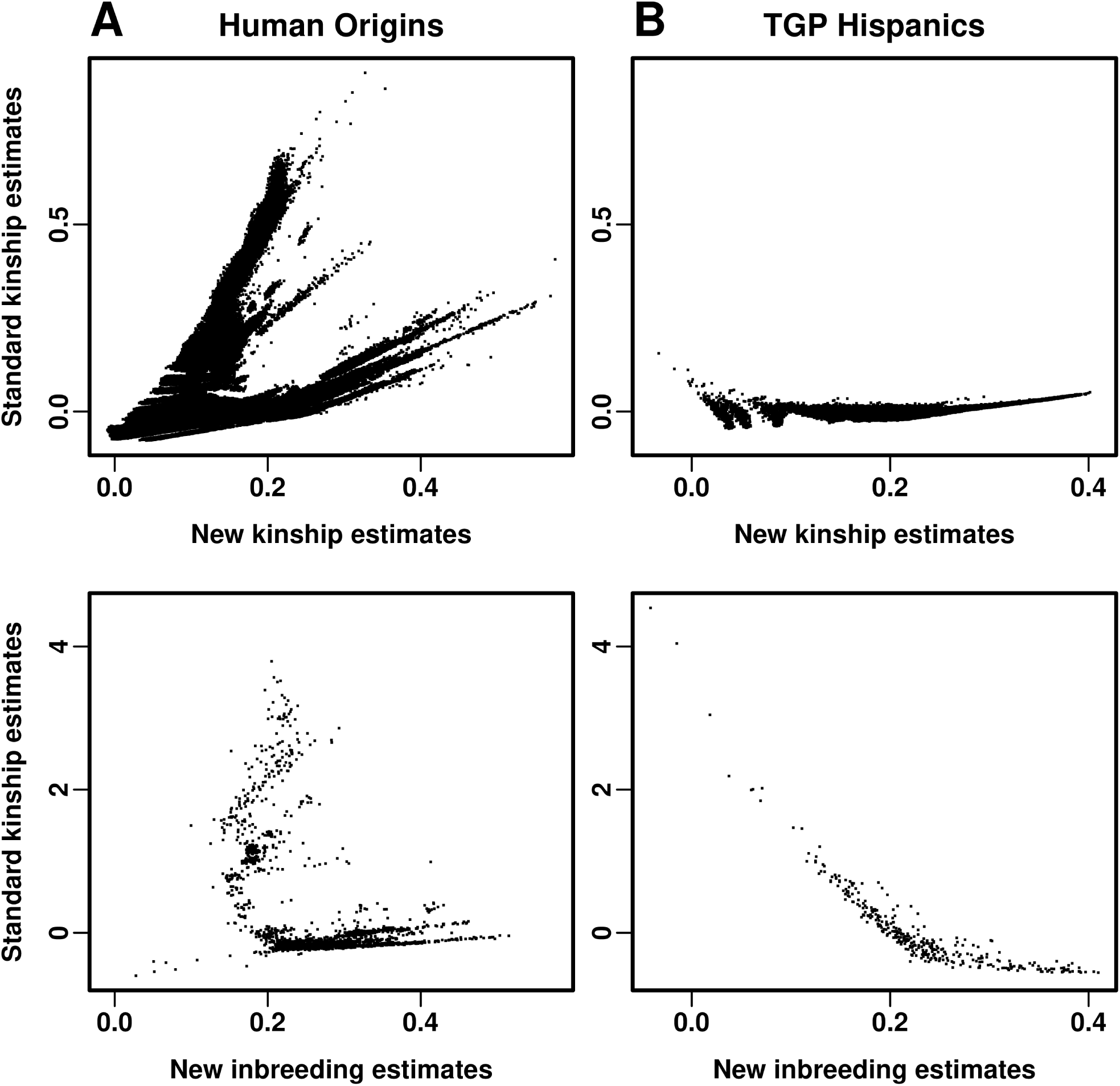
Comparison of new and existing kinship estimates. The x-axes are estimates from the new kinship estimator, while the y-axes are estimates from the standard kinship estimator. Inbreeding coefficients (second row) are compared separately from kinship coefficients (between different individuals; first row) since the scales are very different for the standard kinship estimator (but not for the new kinship estimator). Columns: **A.** Comparison of estimates in the Human Origins dataset. **B.** Comparison of estimates in the 1000 Genomes Hispanics dataset.

**Figure S3:**
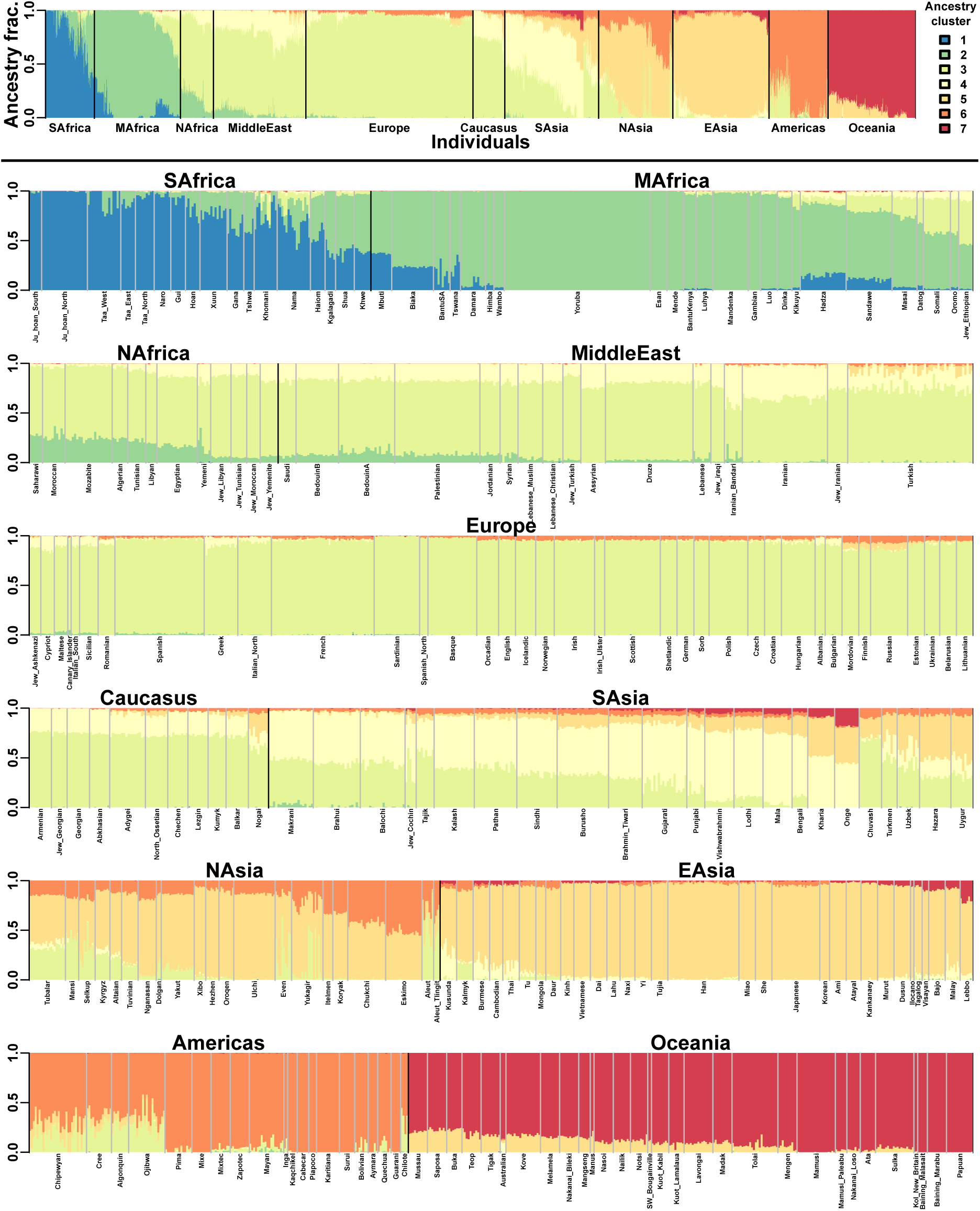
Admixture analysis of Human Origins with *K* = 7. The top row shows the full set of admixture proportions for the *K* = 7 infered ancestry clusters and all 11 subpopulations in Human Origins. All other rows show the same data with greater detail, including labels for every sub-subpopulation. The seven ancestry clusters were ordered manually to correspond roughly with distance from southern Africa.

## S2 1000 Genomes Project

### S2.1 Data processing

The 1000 Genomes Project (TGP) “Phase 3” integrated call data [50, 51] is available at http://www.internationalgenome.org/data (dated 2013-05-02). We started from the plink2-formatted version available at https://www.cog-genomics.org/plink/2.0/resources#1kg_phase3. This dataset was processed using the plink2 software [61]. Our analysis was restricted to autosomal biallelic SNP loci ascertained in `YRI`, after removing loci with repeated identifiers (20,417,484 loci). Of these, 14,145,583 loci are polymorphic in Hispanics (`PUR, CLM, PEL, MXL`; Table S2).

**Table S2:** Overview of 1000 Genomes (TGP) Hispanics dataset.

| Dataset | Loci ($m$) | Individuals ($n$) | Subpopulations |
| --- | --- | --- | --- |
| Full TGP (ascertained in YRI, other locus filters) | 20,417,484 | 2504 | 26 |
| Hispanics | 14,145,583 | 347 | 4 |
| Hispanics + Admixture Panels | 6,216,713 | 665 | 7 |

### S2.2 Admixture analysis

The Admixture analysis of the Hispanic individuals was performed with the addition of individuals from the `YRI, IBS`, and `CHB` subpopulations to help anchor the *K* = 3 admixture clusters. Only loci with minor allele frequency *≥* 10% across the 7 subpopulations (6,216,713 loci, see Table S2) were input to the Admixture software [62]. The cluster associated with `YRI` was assigned to Sub-Saharan African (AFR) ancestry, `IBS` to European (EUR) ancestry, and `CHB` to Native American (AMR) ancestry by proxy (Fig. S4). There are no Native American subpopulations in 1000 Genomes, but the high AMR ancestry predicted for many `PEL` and `MXL` individuals suggests that AMR ancestry is not being underestimated by this procedure.

**Figure S4:**
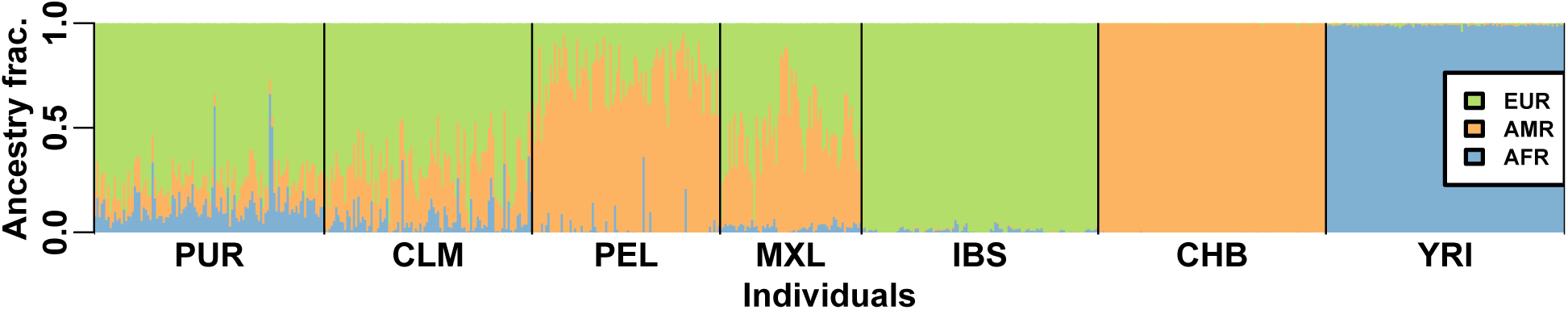
Ancestry inference in Hispanic individuals. Admixture proportions of every individual using YRI, IBS, and CHB as reference panels for Sub-Saharan African (AFR), European (EUR), and Native American (AMR) ancestry, respectively.

### S2.3 Estimation of minimum kinship

The kinship matrix of the Hispanic individuals was estimated as follows. First, the *A*_*jk*_ values were estimated, and the function `seriate` from the R package `seriation` was used with default values to reorder the columns and rows of this matrix so that the lowest kinship values are pushed away from the diagonal [59, 60]. We inspected the individuals at the extremes of the resulting ordering, and found that four individuals with among the highest African admixture proportions also shared the smallest kinship estimates in the data. The two most extreme clusters, (`HG01108, HG01242`) and (`HG01551, HG01241`), were used to estimate *A*_min_, which yields the final kinship estimates 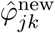.

